# Antimicrobial activity and degradation of superhydrophobic magnesium substrates in bacterial media

**DOI:** 10.1101/2021.04.29.441950

**Authors:** Alexandre M. Emelyanenko, Valery V. Kaminsky, Ivan S. Pytskii, Kirill A. Emelyanenko, Alexander G. Domantovsky, Elizaveta V. Chulkova, Andrei A. Shiryaev, Andrei V. Aleshkin, Ludmila B. Boinovich

## Abstract

The interest in magnesium-based materials is promoted by their biocompatibility, bioresorbability, and by their recently found antibacterial potential. Until now the widespread use of magnesium alloys in different corrosive environments was inhibited by their weakly controllable degradation rate and poorly understood microbiologically induced corrosion behavior. To better understand the degradation and usability of magnesium-based alloys, in this study we have fabricated the superhydrophobic coatings on top of magnesium-based alloy and analyzed the behavior of this alloy in bacterial dispersions of *Pseudomonas aeruginosa* and *Klebsiella pneumoniae* cells in phosphate buffered saline. It was shown that immersion of such coatings into bacterial dispersions causes notable changes in the morphology of the samples, dependent on the bacterial dispersion composition and the type of bacterial strain. The interaction of superhydrophobic coatings with the bacterial dispersion caused the formation of biofilms and sodium polyphosphate films, which provided enhanced barrier properties for magnesium dissolution and hence for dispersion medium alkalization, eventually leading to inhibition of magnesium substrate degradation. Electrochemical data obtained for superhydrophobic samples continuously contacted with the corrosive bacterial dispersions during 48 h indicated a high level of anti-corrosion protection.

## 1. Introduction

In recent years, the interest of the scientific community in the studies of the behavior of magnesium and its alloys in various environments and in the development of new materials based on magnesium for biomedical purposes has sharply increased. This interest is promoted by the biocompatibility of magnesium, the possibility of its use for biodegradable devices, in implantology, and, finally, by the attractive mechanical properties of magnesium alloys. Additionally, the very demanded property of magnesium-based materials is their antibacterial potential, since infections are one of the most unfavorable complications associated with the implantation of any device. At the same time, until recently the widespread use of magnesium alloys in different corrosive environments including the physiological liquids was inhibited by its high degradation leading to the release of hydrogen and an increase in the pH of the liquid medium. In addition, microbiologically induced corrosion of metals usually contributes to the destruction. The rapid and weakly controllable degradation rate of magnesium and its alloys greatly hampers the clinical use of these materials. Therefore, significant efforts are currently targeted on finding the ways to slow down the rate of degradation. Among the most promising options are the use of particles of magnesium or magnesium alloys as fillers for polymer matrices [1–3] and the use of treatments of magnesium alloys, leading to a superhydrophobic state of the surface [4–11].

Recently, we have shown [8] that the appropriate selection of laser treatment regime for obtaining the desired texture and chemical composition of the surface layer makes it possible to create a superhydrophobic coating on a magnesium alloy that exhibits satisfactory corrosion resistance even after prolonged immersion in a 0.5 M sodium chloride solution. In this work, we investigate the degradation and anticorrosive properties of fabricated superhydrophobic coatings under conditions of microbiological corrosion when immersed in bacterial dispersions of bacteria cells *Pseudomonas aeruginosa* and *Klebsiella pneumoniae* in phosphate buffered saline (PBS).

## 2. Materials and methods

### 2.1. Sample preparation

We have used magnesium alloy MA8 with the chemical composition (in weight %): Mn 1.3 – 2.2, Ce 0.15 – 0.35, impurities <0.3, Mg balance, to fabricate the superhydrophobic samples for studies described in this paper. Flat sheets with a thickness of 2 mm were cut on samples with a size of 20×20 mm^2^. Surface texturing was performed using nanosecond laser treatment with laser processing regimes described in our recent paper [8]. The selection of the regimes was based on the best corrosion protection properties of samples during prolonged contact with 0.5M NaCl aqueous solution. To perform the laser processing of the samples, we used an Argent-M laser system (Russia) with an IR ytterbium fiber laser (wavelength of 1.064 μm, nominal power 20 W, a beam waist of 40 μm), and a RAYLASE MS10 2-axis laser beam deflection unit (Germany). The laser beam used for the raster scanning of the sample was characterized by the following parameters: fluence 1.5 J/cm^2^, line density 400 mm^-1^, linear scanning rate 100 mm/s, pulse duration 4ns, repetition rate 1000 kHz. Laser treatment was performed at a temperature of 20-25 °C and relative humidity of 40-50%. After the laser treatment, the samples were thoroughly washed with deionized water to remove surface micro-and nanoparticles weakly adhered to the sample.

To attain the superhydrophobic state of the surface, the laser-treated samples were first subjected for 1 h to ozone-assisted UV irradiation, and then hydrophobized by exposure to saturated vapors of methoxy-{3-[(2,2,3,3,4,4,5,5,6,6,7,7,8,8,8-pentadecafluorooctyl)oxy]-propyl}-silane in the sealed vessel for 1 h at 105 °C and dried for 1 h in an oven at 150 °C. The details of the hydrophobization procedure with substantiation of each stage necessary to get the superhydrophobic state with extremely high contact angle, low roll-off angle, and satisfactory durability were discussed in [12].

### 2.2. Analysis of bactericidal properties of the superhydrophobic magnesium substrates

In this study, the bactericidal activity of superhydrophobic magnesium alloy substrates was studied with respect to two pathogenic strains, *Pseudomonas aeruginosa* (*P. Aeruginosa*) and *Klebsiella pneumoniae* B-811 (*K. pneumoniae*), isolated from hospital patients. Phosphate buffered saline (PBS) purchased from VWR Life Science AMRESCO (USA) was used as a dispersion medium. To prepare the bacterial suspensions, an overnight bacterial culture was introduced into the PBS and incubated for 18 h at 37 °C. For the experiments, dispersions of bacterial cells with an initial titer of 10^7^ – 10^8^ colony-forming units (CFU) per mL were used.

To test the bactericidal properties with respect to planktonic forms of different strains, superhydrophobic plates were placed in a separate sterile cup for each strain, and 40 mL of bacterial suspension was poured into the cup. Cups were stored at room temperature for 48 h. To evaluate the bactericidal action of the plates in contact with the dispersion, after a predefined time of contact, an assay of a bacterial suspension with a volume of 0.5 ml was taken from each cup. Ten-fold dilutions were prepared, then 0.1 mL was taken from each dilution and evenly distributed over the surface of Müller-Hinton agar in Petri dishes. After the incubation at 37 °C for 24 h, - the bacterial titer was determined by counting the formed colonies. To quantify the effect of contact between the bacterial suspension and the superhydrophobic MA8 plate, for each strain and each contact duration the obtained titer was normalized to the initial titer in the dispersion.

Additionally, for each strain and each contact duration, the second assay (1 mL) was used for the determination of magnesium egress into the PBS, as described below.

### 2.3. Determination of the concentration of Mg^2+^ in a dispersion medium

The assay with 1mL of the bacterial dispersion was poured into an Eppendorf tube and centrifuged at 14000 rpm for 10 min. The liquid phase after centrifugation was used to determine magnesium content in the PBS by mass spectroscopy analysis, using an inductively coupled plasma mass spectrometer (ICP-MS) Agilent 7500CE (Agilent Technologies, USA).

Concentrated nitric acid (65 wt.%) was added to each sample for obtaining 0.01 M solution. Then, the digestion of samples was carried out at 60 °C for 40 min. Nitric acid used in the digestion was high-purity-grade (Sigma-Aldrich). All digest samples were prepared in a laminar flow cabinet. The resulting solution was injected directly into the instrument. Laboratory blank samples were also analyzed for the same digestion method. The average magnesium concentration in blank samples was then subtracted from the measured concentration in each digest to give the final reported concentration of that element. Each sample was analyzed 3 times in order to assess possible drift effects. Standard solutions for calibration with respect to magnesium were prepared by diluting multi-element stock solutions (Agilent Technologies multi-element solution, 10 mg/L) with Milli-Q water containing 0.6% (0.01 M) nitric acid. Calibration scale was from 1 μg/L to 1 mg/L, R^2^ = 0.999.

### 2.4. Characterization of surface morphology, chemical and phase composition

Phase analysis of laser-treated samples was performed with X-ray diffractometer Empyrean (Panalytical BV, The Netherlands) using Ni-filtered Cu Kɑ_1_-radiation in standard Bragg-Brentano (reflection) geometry. Diffraction patterns of the as-treated samples were dominated by very strong reflections from bulk Mg-alloy; identification of the phases in the surface layer was complicated both by a very small amount of material and plausible texture effects, making interpretation difficult. Subsequently, material from the surface was collected by gentle mechanical scratching, thus minimizing the contribution of the bulk material and decreasing the texture influence.

The morphology and the elemental composition of samples were studied by field-emission scanning electron microscopy and energy-dispersive X-ray spectroscopy (EDS) on a FIB-SEM Nvision 40 workstation (Zeiss, Germany) equipped with X-MAX energy-dispersive detector (Oxford Instruments, UK). The SEM images were recorded in secondary electron detection mode at accelerating voltages of 2–5 kV. EDS microanalysis was performed at 10 kV accelerating voltage.

The infrared spectra of the samples were investigated using Fourier-transform infrared (FTIR) spectrometer Nicolet 6700 (Thermo Scientific, USA) using specular apertured grazing angle (SAGA) accessory and a mercury cadmium telluride (MCT) detector cooled with liquid N_2_. The angle of incidence for SAGA was 80 degrees, and the diameter of the circular sampling area was 8 mm. The spectra were recorded at a resolution of 4 cm^−1^. All the spectra were derived from the result of an average of 128 scans.

### 2.5. Characterization of surface wettability and pH of the dispersion medium

The measurement of pH in bacterial dispersions after a predefined time of contact between the superhydrophobic sample and the dispersion was performed using microelectrode ESK-10614 (LLC Measuring Technology, Russia).

The wettability of the coatings just before and after contact with bacterial dispersions was characterized by measuring the contact and roll-off angles. Digital image processing of sessile droplet and Laplace fit optimization for determining the droplet shape parameters were used to calculate the contact angles [13]. The roll-off angle for the sessile droplets was defined upon smooth substrate tilting until the droplet started to roll over the surface. Both the contact angles and roll-off angles were measured for 15 μL droplets at least at 10 different surface locations for each sample.

### 2.6. Characterization of electrochemical properties of surfaces after contact with bacterial suspensions

The electrochemical properties of the test surfaces were studied using an electrochemical workstation Elins P50x (Elins, Russia) equipped with a frequency-response analyzer FRA 24M for electrochemical impedance spectroscopy measurements. The measurements were carried out at 25 °C in a three-electrode cell PAR K0235 (Princeton Applied Research, USA) with a 0.5 M NaCl aqueous solution as an electrolyte. A silver/silver chloride electrode (Ag/AgCl) filled with saturated KCl solution served as a reference electrode and a Pt mesh as a counter electrode.

To study the effect of corrosion degradation of superhydrophobic samples in bacterial dispersions, the samples were immersed in the dispersion for a certain time up to 48 h. The corrosion degradation was analyzed through the corrosion current and the impedance spectra. For the measurements, the sample was taken off the dispersion, rinsed with a distilled water, and placed into the three-electrode electrochemical cell as a working electrode. The measurement of polarization curves was performed after 60 min equilibration of the sample to a 0.5M sodium chloride aqueous solution. The potentiodynamic polarization curves were registered at a scan rate of 1 mV/s in the applied potential range from −150 to +300 mV (with respect to open circuit potential). The corrosion potential, *E_cor_*, and current, *I_cor_*, were derived from the potentiodynamic polarization curves after Tafel extrapolation.

A sinusoidal perturbation signal with an amplitude of 10 mV (with respect to open circuit potential) was used for the electrochemical impedance spectroscopy measurements. Impedance spectra were acquired in the frequency range from 0.05 Hz to 100 kHz with a logarithmic sweep (20 points per decade).

## 3. Morphology and wettability of superhydrophobic coating

The as-fabricated superhydrophobic coating on magnesium alloy MA8 is characterized by the hierarchical roughness and low surface energy which resulted in an apparent contact angle of 171±1° and roll-off angle of 3±1°. The morphology of the textured surface of this coating before contact with corrosive liquid medium is shown in Figure 1a,b. Immersion of this coating into the bacterial dispersions in phosphate buffered saline (PBS) causes notable morphology changes (Figure 1c-h) because of the high corrosive activity of such liquid medium.

**Figure 1.**
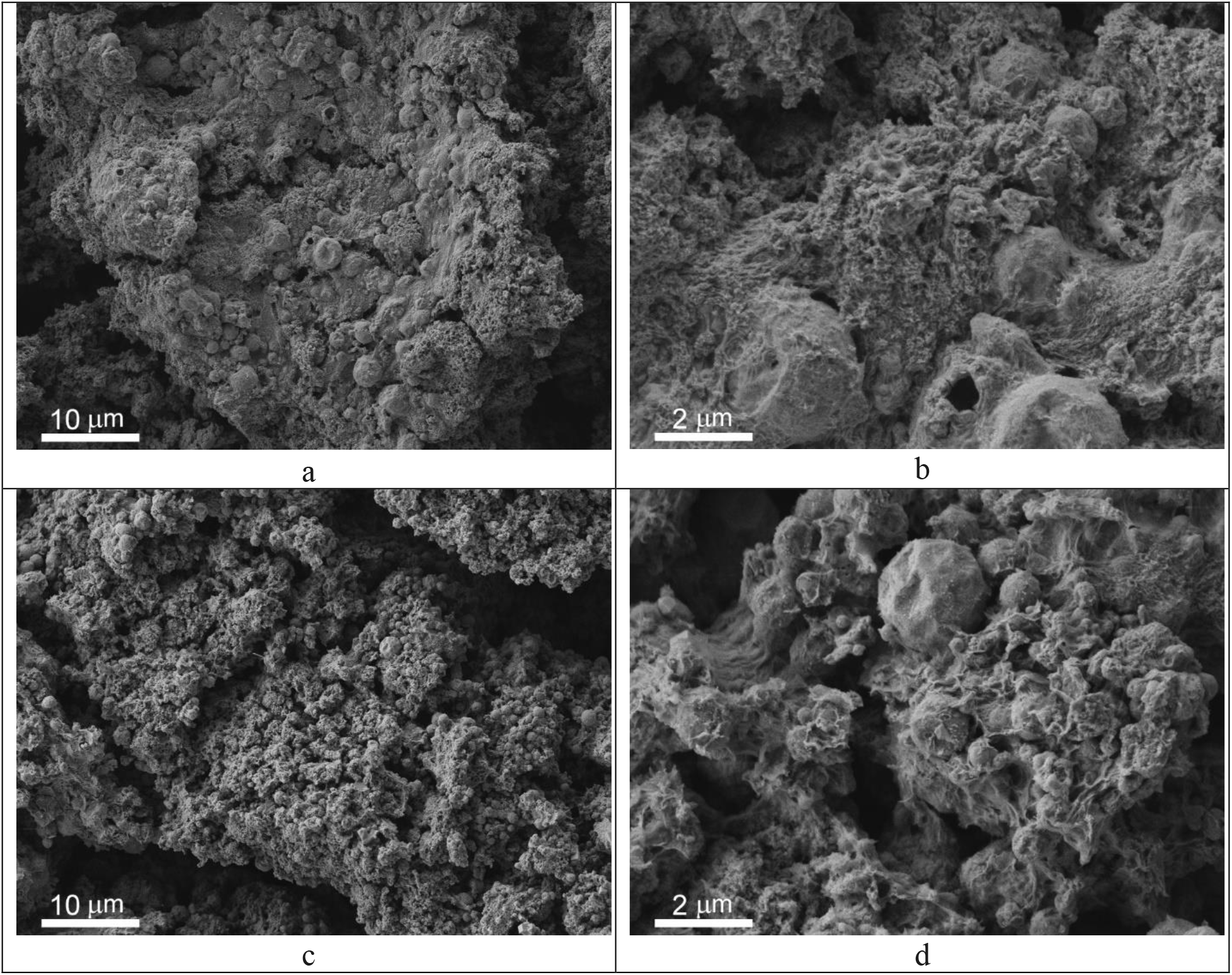
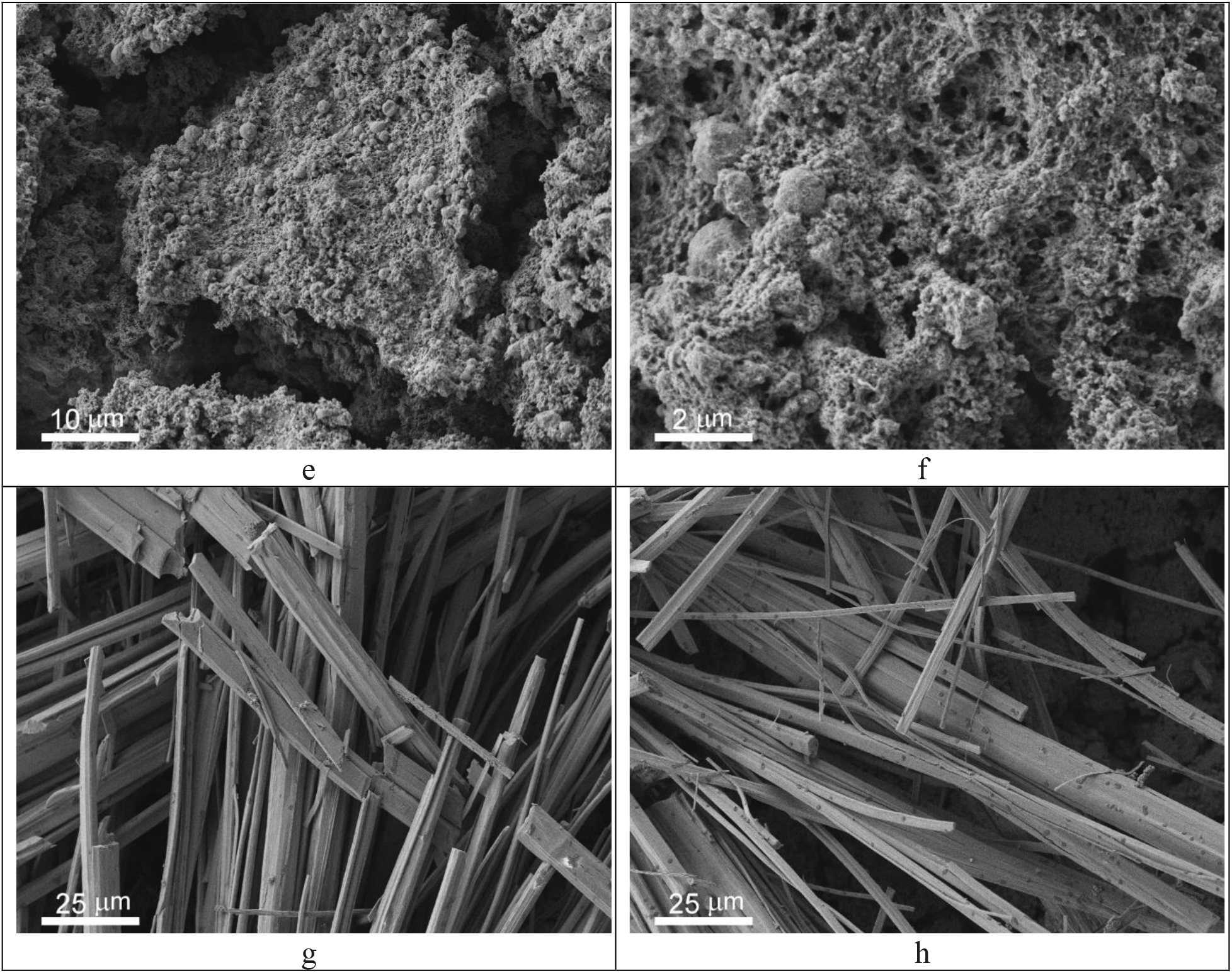
SEM images illustrating the surface morphology of superhydrophobic coating on MA8 magnesium alloy: (a,b) freshly prepared; (c,d) covering film after 48 h of immersion into bacterial dispersions of *K. pneumoniae* in phosphate buffered saline (PBS); (e,f) covering film after 48 h of immersion into bacterial dispersions of *P. aeruginosa* in PBS, (g) microrods formed during 48 h of immersion into bacterial dispersions of *K. pneumoniae* in PBS; (h) microrods formed during 48 h of immersion into bacterial dispersions of *P. aeruginosa* in PBS.

The morphology of superhydrophobic MA8 surface after 48 h of contact with *P. Aeruginosa* and *K. pneumoniae* has some similar features, like bundles of microrods, formed as a result of the interaction of magnesium with phosphate buffered saline (Figure 1 g,h). However, the newly formed surface layer covering the intrinsic features of the superhydrophobic surface, although preserves the multimodal surface texture, differs from the original surface and is different for the samples contacted to different bacterial dispersions (compare Figure 1 b, d, and f).

It is well documented in the literature that bare magnesium and its alloys are characterized by high intrinsic corrosion propensity in the aqueous phase. Anodic and cathodic electrochemical reactions of magnesium in an aqueous solution are presented by the equations

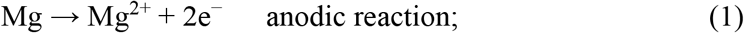

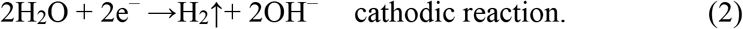

The whole electrochemical corrosion process is described by the reaction

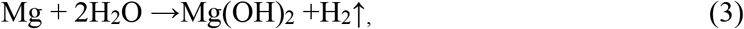

which can be easily detected through the hydrogen release and alkalization of the liquid phase. Besides, an additional indication of the corrosive process at pH>10.5 is the precipitation of magnesium hydroxide [14]. The precipitated magnesium hydroxide forms a layer on the magnesium surface; however, this layer does not provide protection of the surface against further magnesium dissolution. The replacement of water by electrolyte with complex composition, for example, by biological liquids modifies the corrosion process and diversifies the corrosion products, kinetics, type, and degree of surface degradation [14, 15]. The presence of bacterial cells in the liquid medium, contacting the metal surface can either enhance the corrosion process due to combination with microbiologically induced corrosion or suppress corrosion degradation [16, 17]. The morphology of the superhydrophobic MA8 surface after 48 h of contact with dispersions of *P. Aeruginosa* and *K. pneumoniae* has some similar features, like bundles of microrods, formed as a result of the interaction of magnesium with phosphate buffered saline. However, the newly formed surface layer covering the intrinsic features of the superhydrophobic surface, although preserves the multimodal surface texture, differs from the original surface and is very different for the sample contacted to different bacterial dispersions (compare Figure 1 b, d, and f). To analyze the degradation of magnesium alloy MA8 with a superhydrophobic coating during immersion in PBS or bacterial dispersions, we have studied the variation of different properties of both the substrates and the liquid medium.

Let us first consider the variation of the superhydrophobic sample wettability during 48 h of contact with liquid medium. It was found that just after withdrawal of the sample from the cup with PBS free of bacterial cells, the wettability was notably different from that obtained for the sample before contact with the corrosive medium. The measured contact and roll-off angles became 163.3±1.0° and 14.3±3.9°, respectively. However, washing with water jet followed by 30 min of drying in an oven at T=180 °C led to nearly complete recovery of the superhydrophobic state with a contact angle of 170.7±1.0° and roll-off angle of 3.5±2.0°. The analysis of the variation in the wettability of superhydrophobic samples during the contact with bacterial dispersion was performed for samples, subjected to immersion into bacterial dispersions, followed by immediate washing by water jet and drying in an oven at T=180 °C for 30 min. Such immediate heat treatment after sample withdrawal from the dispersion was performed to ensure complete bacterial decontamination. The measurement of the wettability of the sample after described procedure indicated preservation of the superhydrophobic state for all studied samples in both bacterial strain dispersions. As an example, Figure 2 shows the values of contact and roll-off angles for one set of experiments with the dispersion of *K. pneumoniae* at immersion time duration from 1 to 48 h. Two values are given for each immersion time, one corresponding to the sample before immersion (blue symbols) and the other after a certain time of immersion (red symbols).

**Figure 2.**
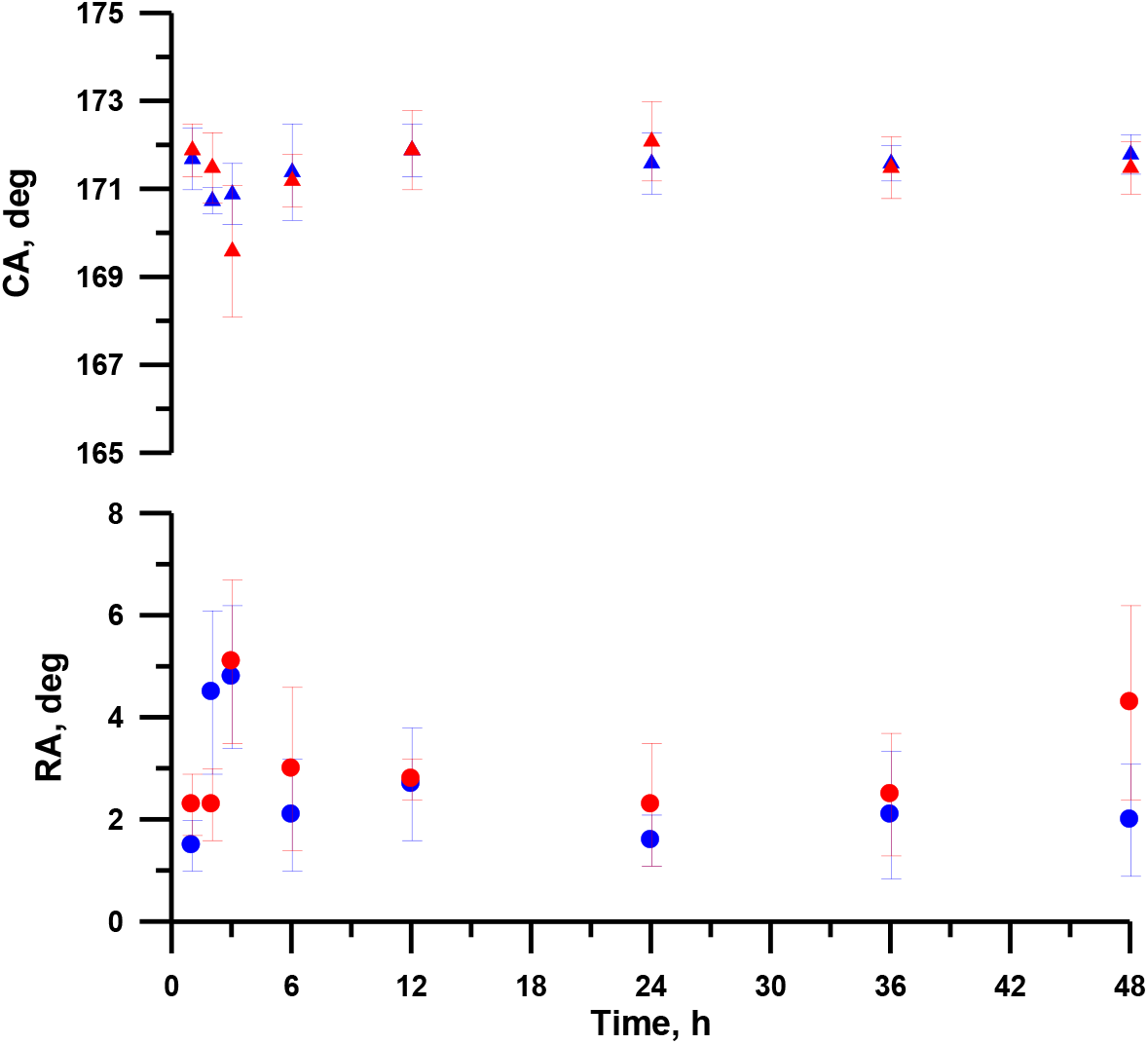
The contact (triangles) and roll-off (circles) angles for the superhydrophobic samples before (blue symbols) and after (red symbols) different time of immersion into the dispersion of *K. pneumoniae*. A separate freshly prepared sample was used for each immersion time, that is why the initial values (blue symbols) show some spread.

As it can be concluded from the analysis of presented data, the degradation of a superhydrophobic state for all samples is weak, if any. At the same time, the significant difference in the morphology allows suggesting the modification of the surface layer composition during contact with the bacterial dispersions. That is why the second step in our study was related to the analysis of the surface composition.

## 4. Variation in surface composition of a superhydrophobic coating

X-ray diffraction patterns (Figure 3a) of the superhydrophobic samples, one of which (1) did not contact to corrosive medium, and two others (2 and 3) were immersed for 48 h into bacterial dispersion of *K. pneumoniae* or *P. Aeruginosa* respectively, are dominated by reflections from MgO. Such oxide is formed in the course of interaction of laser irradiation with the magnesium alloy surface in the presence of atmospheric oxygen. In samples 2 and 3, the amorphous material is abundant. Unambiguous phase identification of other surface phases is difficult both due to rich variations in cation composition, degree of crystallinity and hydration of many complex phosphates. Orthorhombic hydrated sodium polyphosphate Na_6_P_6_O_18_·6H_2_O is observed on sample 2 and in a smaller amount on sample 3. Weak reflections from other crystalline phases were also found, but their assignment is not univocal. Possible candidates include varieties of Mg, Na, K hydrophosphates.

**Figure 3.**
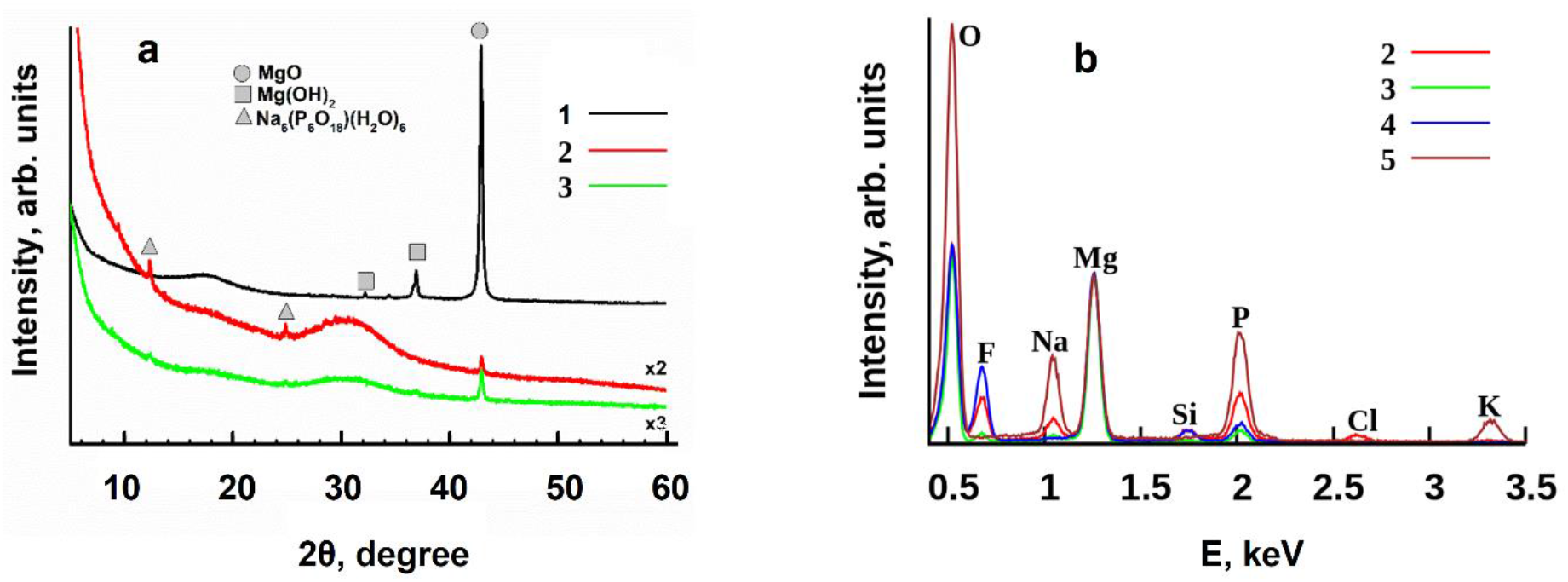
X-ray diffraction patterns (a) and EDS spectra (b) for the superhydrophobic samples before contact with biological liquids (1) and after 48 h of immersion into dispersions of *K. pneumoniae* (2), *P. Aeruginosa* (3), or PBS without bacteria (4). EDS spectrum (5) was measured from bundles of micro rods (like those in Figure 1 a,c,e) and was characteristic of micro rods formed in both PBS dispersions studied, as well in PBS without bacteria.

EDS spectra (Figure 3b) were measured separately for the rods and the areas free of micro rods, grown on the surface of samples 2 and 3 in the bacterial dispersions. It was found that the rod’s composition was not sensitive to the type of the bacterial cells and was the same for the two bacterial dispersions in PBS and for PBS free from the cells. The main components of the rods are Mg, Na, K hydrophosphates. Covering films formed on top of superhydrophobic surfaces in two studied bacterial dispersions differ from each other by the amount of phosphorous, potassium, and sodium. Besides, notably lower height of the peaks corresponding to the components of the very top layer of the superhydrophobic texture, such as silicon and fluorine, was registered by EDS. Jointly with the data of X-ray diffraction, indicating the presence of the amorphous material, a decrease in the height of F and Si peaks at the preservation of a superhydrophobic state of the sample indicates the presence of hydrocarbon deposits. Such deposits formed on the surface as a constituent of biofilm, resulting from the interaction of bacterial cells with the superhydrophobic surface. The thicker the newly formed biofilm, the weaker the EDS signal from the superhydrophobic coating. To examine this hypothesis, we have measured FT-IR reflectance spectra of the surface in the range of wave numbers 3500-2500 cm^-1^, corresponding to stretching C-H vibrations (Figure 4). Indeed, higher C-H absorption leading to smaller reflectance of IR irradiation from the surface was detected for the sample contacted to *P. Aeruginosa* cells in comparison to the sample immersed into *K. pneumoniae* dispersion. Such behavior indicating thicker biofilm in the former case well correlated with lower height of F and Si peaks in EDS spectra.

**Figure 4.**
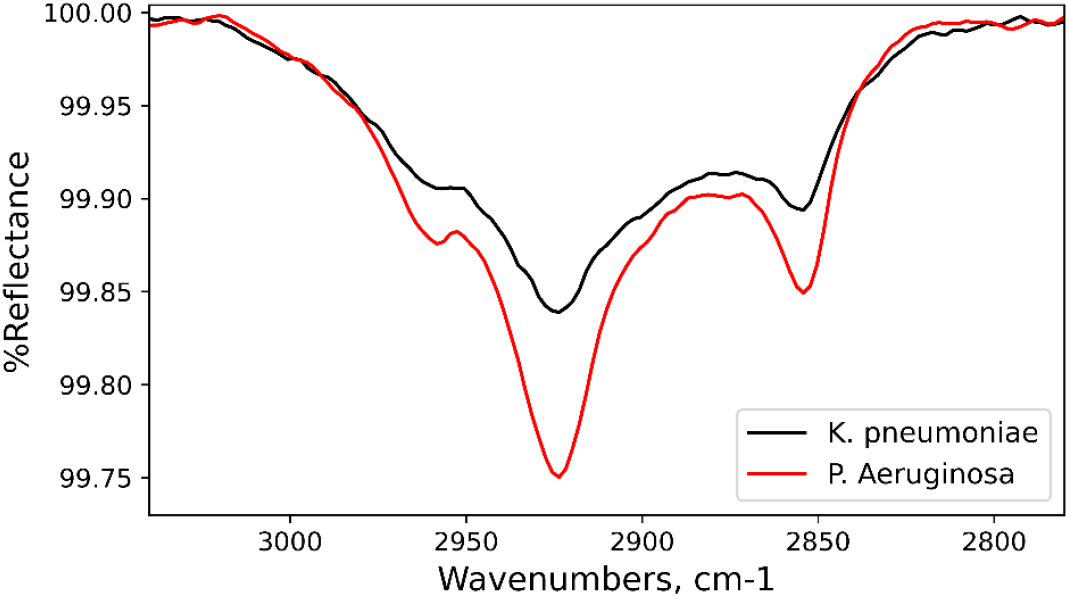
FT-IR reflectance spectra for the surface of superhydrophobic samples after 48 h of immersion into dispersions of *K. pneumoniae* and *P. Aeruginosa*.

The formation of such a biofilm on top of a superhydrophobic surface should affect the bactericidal action of magnesium with respect to the bacterial cell, the alkalinity of the liquid phase, and the corrosion resistance of the superhydrophobic coating during the contact with bacterial dispersions.

## 5. Antibacterial activity of superhydrophobic coatings in bacterial dispersions

The bacterial activity of superhydrophobic samples of MA8 alloy immersed into bacterial dispersion with respect to planktonic forms of *K. pneumoniae* or *P. Aeruginosa* was estimated by monitoring the evolution of bacterial titer in the dispersion with the immersed sample. The analysis of obtained data presented in Figure 5 indicates that the titer of *K. pneumoniae* started to notably decrease only after 12 h of contact between the bacterial dispersion and a metal sample with the superhydrophobic coating. Such depression of antibacterial activity of metal plate compared to bare metal is related to the high protective action of a superhydrophobic surface, which inhibits the transition of ions, charges, and water molecules through the superhydrophobic textured layer, suppresses the adhesion of bacterial cells onto the surface, and significantly decreases the rate of metal dissolution. However, after 48 h, the dispersion can be considered free from bacterial contamination. Since the wettability studies described above indicated the preservation of a superhydrophobic state of the sample after 48 h of contact between the sample and the dispersion and had shown only an insignificant decrease in contact angles and an increase in roll-off angles, the observed bactericidal action is seemingly related to the appearance of a few wetting defects.

**Figure 5.**
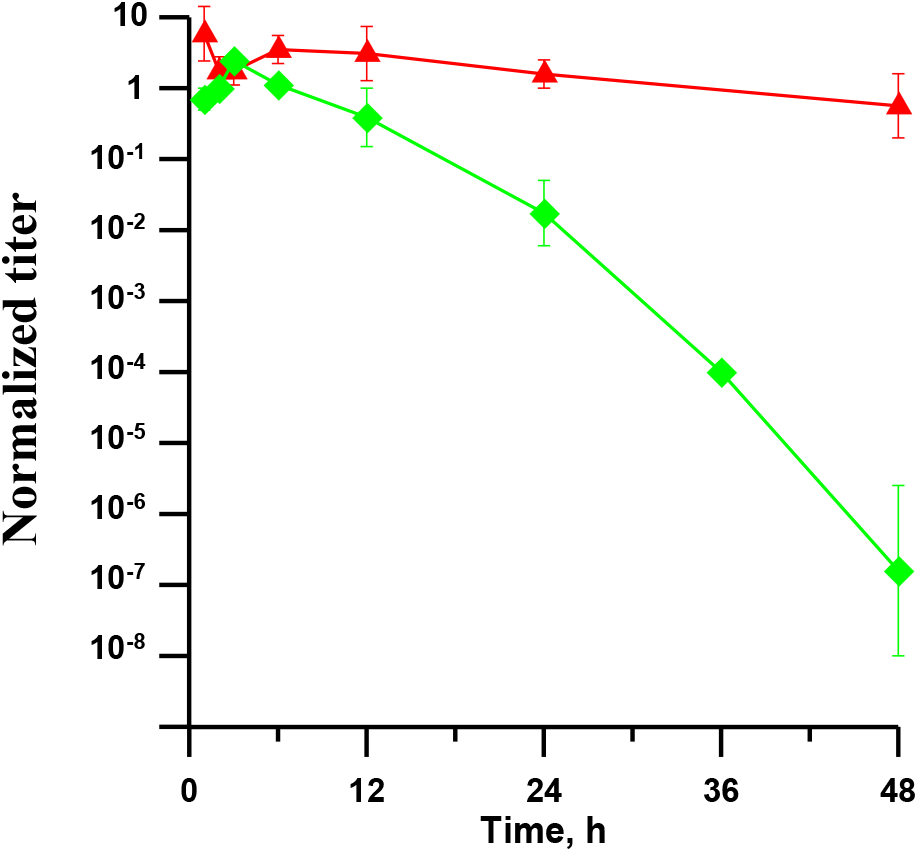
Evolution of bacterial titer of *K. pneumoniae* (green diamonds) and *P. Aeruginosa* (red triangles) during prolonged contact between the corresponding bacterial dispersion and immersed MA8 alloy sample with the superhydrophobic coating. The normalized titer was calculated as a ratio of the titer in an assay taken at a given time to the initial titer in the dispersion.

In contrast, the biofilm formed on top of the superhydrophobic surface immersed in bacterial dispersion of *P. Aeruginosa* additionally inhibited the bactericidal action of the metal. Even after 48 h of contact with the superhydrophobic sample, the titer of *P. Aeruginosa* cells in the dispersion remained as high as 50% of the initial titer.

Analysis of the literature shows that there is still no consensus on the nature of the anti-bacterial activity of magnesium alloys. The main mechanisms of toxic action discussed in the literature are [14]: (1) high reactivity of Mg in contact with aqueous media, leading to the formation of superoxide ions (O^2-^); (2) an excess of magnesium ions in the aqueous medium surrounding the cells, leading to osmotic effects that destroy cell’s membranes; (3) an increase in pH during corrosion of magnesium in biological media. Additionally, two mechanisms specific for any superhydrophobic material should be taken into account [18–21]: (4) low adhesion of bacterial cells to the superhydrophobic surface; and (5) mechanical damage of cell membranes for the cells deposited onto the surface. The analysis of mechanism (1) related to oxidation stress is out of the scope and technical capabilities of this study. The antibacterial mechanisms (4) and (5) related to the superhydrophobic state are significant, but not determinative for the planktonic forms of bacteria. The evolution of pH in PBS and bacterial dispersions, as well as the behavior of Mg^2+^ ions concentration in dispersion medium we will discuss in the next session.

## 6. Variation of pH in dispersion medium during its contact with a superhydrophobic sample

To monitor the variation of pH, which can be considered as an indicator of magnesium alloy dissolution in both PBS, free from bacterial cells, and in bacterial dispersions in PBS, the magnesium alloy samples were immersed into the liquid with the ratio of the apparent sample surface to the liquid volume equal to 0.8 cm^2^/1 mL, as described above. Data on the variation of pH are presented in Figure 6.

**Figure 6.**
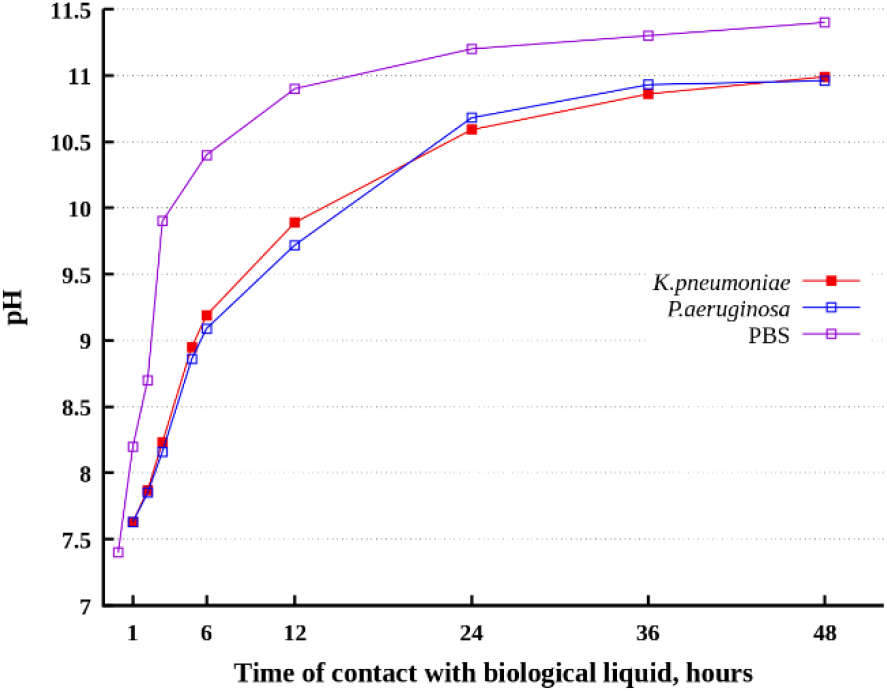
Variation of pH with a time of contact between MA8 alloy sample with the superhydrophobic coating and phosphate buffer solution (PBS) with or without bacterial cells.

After immersing the superhydrophobic sample for 48 h into PBS free from bacterial cells, the rapid increase in pH of liquid from 7.4 to 11.4 was detected. Such alkalization is taking place due to the cathodic reaction (2).

Here it is worth noting that the superhydrophobic state of magnesium surface provides notable anticorrosion protection in neutral chloride-containing media. However, a significant pH increase causes the hydrolysis of Si-O and Si-C bonds in the molecules of fluorooxysilane, used here as a hydrophobic agent [12]. Such hydrolysis results in hydrophobic molecules desorption from the surface and formation of the wetting defects, which act as channels for ions and charges transfer through the surface layer of the sample and thus causing some degradation of magnesium alloy.

In the presence of chloride ions in the alkaline solution, the substitution reaction transforms magnesium hydroxide into soluble chloride, which allows for an increase in pH to values higher than pH=10.5, corresponding to magnesium hydroxide precipitation. In turn, the interaction of magnesium ions with KH_2_PO_4_ monopotassium phosphate or Na_2_HPO_4_ disodium phosphate will cause the formation of weakly soluble mixed magnesium, potassium and sodium phosphates, which deposited onto the surface as covering film (Figure 3b) or rod-like crystals (Figure 1).

Data presented in Figure 6 for the pH evolution in bacterial dispersions indicate partial suppression of alkalization of dispersion medium in the presence of the bacterial cell. The observed phenomena can be related to the additional protective action of covering surface film, which grew on the superhydrophobic surface in bacterial dispersion in PBS. At the same time, even in the presence of bacterial cells, the pH reached values as high as pH=11.0.

The capturing of magnesium by weakly soluble compounds is additionally substantiated by low magnesium concentration in a liquid medium. Having in mind that the obtained concentrations of Mg^2+^ are notably lower than those discussed in the literature as toxic and have an order of the typical concentrations in the cellular liquids [14, 22, 23], we can exclude from consideration the mechanism of bacteria killing associated with the osmotic effects caused by super high concentrations of magnesium in the systems under consideration.

Nonmonotonic increase in Mg^2+^ concentration in time for all three considered liquid media is seemingly determined by the kinetics of formation of the covering films and the rods on the surface of superhydrophobic samples. Since the superhydrophobic surface is characterized by an increased energetic barrier for the heterogeneous nucleation of the surface phases [24], we observe the increase in the concentration of magnesium ion till reaching the components saturation necessary to start the mass formation of a new phase (Figure 7).

**Figure 7.**
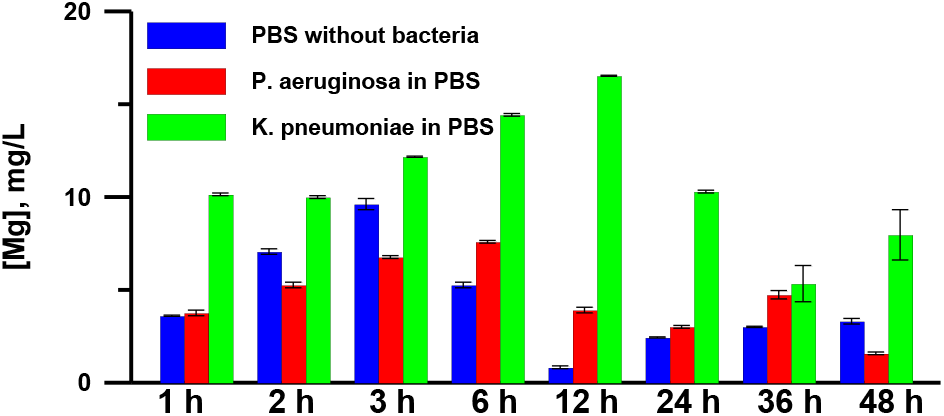
Variation of magnesium concentration in liquid phase with a time of contact between MA8 alloy sample with the superhydrophobic coating and phosphate buffer solution (PBS) with or without bacterial cells.

To present more information on the degradation behavior of superhydrophobic surfaces in PBS and bacterial dispersions in PBS the electrochemical parameters of the superhydrophobic sample will be given in the next section.

## 7. Anticorrosion behavior of superhydrophobic coatings

The superhydrophobic coating on the MA8 surface, obtained by laser treatment followed by chemisorption of fluorooxysilane, works as a good protective anticorrosive coating, inhibiting the evolution of hydrogen and the formation of crystalline hydrates on the surface. The change in the state of such samples and their possible corrosive degradation were quantitatively characterized in section 3; here we will use the methods of polarization curves and impedance spectroscopy for deeper analysis. The corrosion currents and the spectra of the impedance modulus were measured before and after immersion of the samples in the bacterial dispersion for 1, 2, 3, 6, 12, 24, 36, and 48 h. It is worth noting that high-quality superhydrophobic surfaces are characterized by extremely low corrosion currents and high impedance values at low frequencies. That is why even minor wetting defect on the surface may cause an order of magnitude variation of the corrosion current and the modulus of impedance. To avoid uncertainty in the initial characteristics of the samples used for this study, we have compared the corresponding characteristics for each sample before and after contact with bacterial dispersions. Two different samples were used for each time of immersion of sample into bacterial dispersion and for each type of dispersion. Figure 8 shows the obtained values of corrosion currents and impedance moduli before and after contact with the bacterial dispersion for individual superhydrophobic samples immersed in the dispersion for a certain time.

**Figure 8.**
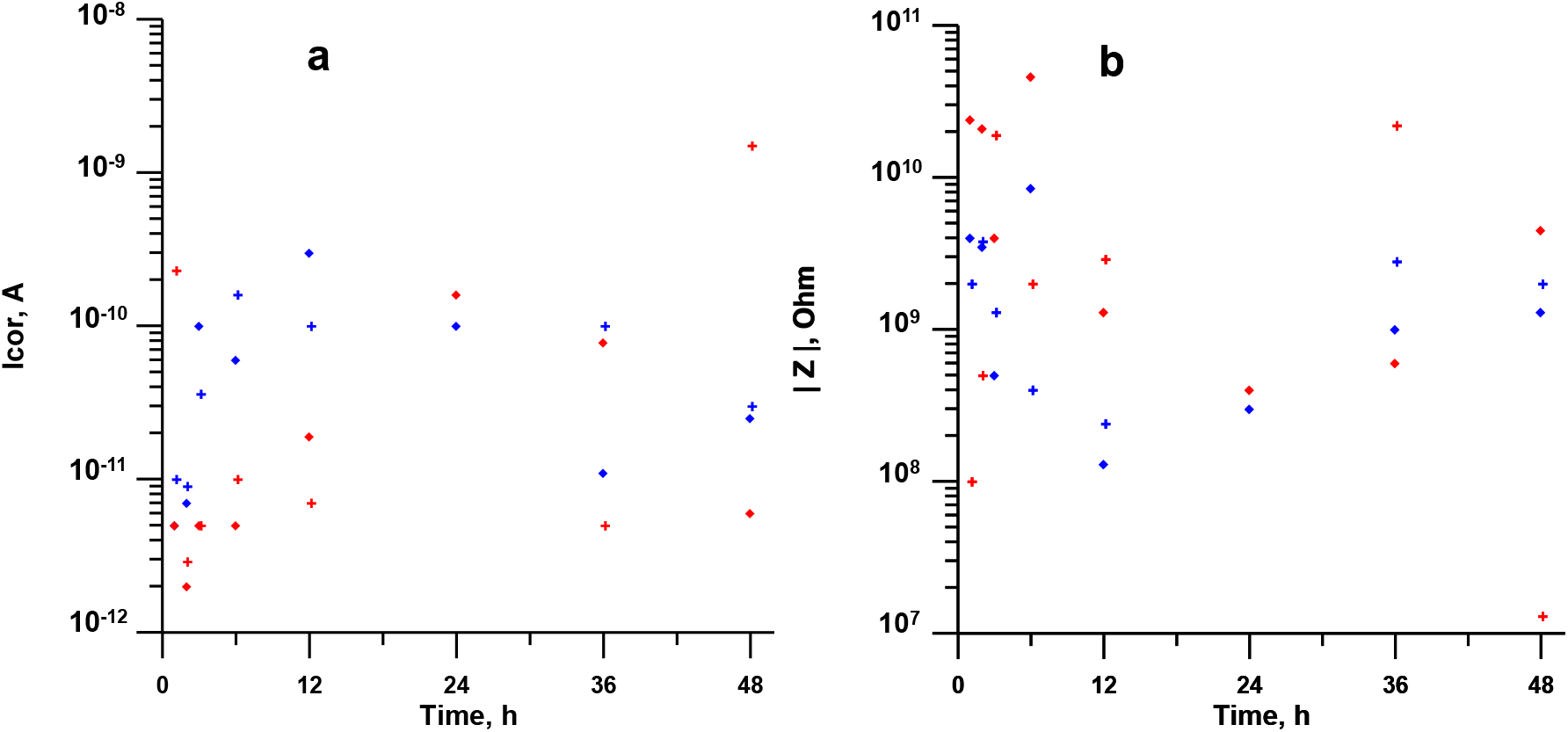
Change of corrosion current (a) and low-frequency (*f*=0.05 Hz) impedance modulus (b) at different times of immersion of superhydrophobic magnesium samples into *P. aeruginosa* dispersion in PBS. Blue and red symbols denote the values measured before and after immersion, respectively; for the given time of immersion, the same symbols (crosses or diamonds) are related to the same sample.

The presented data indicate weak changes in the electrochemical parameters of coatings even with prolonged, for 48 h, contact of the magnesium alloy with a corrosive biological environment. In this case, for a majority of samples no degradation of the anticorrosive properties of the coating was detected, but, on the contrary, an increase in the corrosion resistance was indicated by a decrease in corrosion currents and an increase in impedance moduli. Possible mechanisms of inhibition of biocorrosion on different metal surfaces including superhydrophobic surfaces, discussed in the literature earlier [17, 25, 26], are as follows: (1) suppression of the growth of corrosive bacteria by antimicrobial drugs; (2) release of a peptide corrosion inhibitor; (3) formation of a protective layer in the case of the physiological activity of bacteria or in the case of strong adhesion of bacterial cells to the interface; (4) formation of a protective layer from corrosion products; and, finally, (5) removal of corrosive agents due to the physiological activity of bacteria. Our data allow us to associate corrosion inhibition in the systems considered here with two factors. This is, firstly, the formation of a protective layer due to the physiological activity of bacteria. The formation of such a layer, referred to here as a biofilm and containing hydrocarbon components and mixed phosphates of magnesium, potassium, and sodium, is indicated by the data of IR spectroscopy (Figure 4) and EDS (Figure 3b). The second factor is related to the formation of layers of corrosion products – magnesium phosphates. Besides, XRD data indicate the precipitation of sodium polyphosphate from the liquid phase onto the superhydrophobic surface. Phosphate coatings, such as magnesium phosphate or sodium polyphosphates formed by different methods have been repetitively employed in earlier literature and were shown able to improve the anti-corrosion performance of Mg-based biomaterials [27–31].

## 8. Conclusions

Bare magnesium alloy MA8 is characterized by low corrosion resistance in different media, leading to rapid degradation of magnesium-based materials. It was shown in recent studies that the fabrication of superhydrophobic coatings on the surface of such materials significantly enhances the stability of materials against different types of degradation, including the corrosion processes. In this study, we have analyzed the behavior and the impact of the interaction of superhydrophobic MA8 samples with the different corrosive media. Experiments were performed in phosphate-buffered saline and the dispersions of bacteria cells *P. Aeruginosa* and *K. pneumoniae* in PBS.

The superhydrophobic samples used were characterized by extremely high initial corrosion resistance in chloride ions solutions. Immersion of such samples into bacterial dispersions resulted in notable changes in the morphology of the sample, due to deposition of the corrosion products related to the formation of a covering film and bundles of rods of magnesium phosphates and mixed magnesium, potassium, and sodium phosphates. Corrosion reactions caused a significant rise in pH and consequently, the bacterial cells killing. It was found that the bactericidal activity of superhydrophobic coatings is dependent on the type of the bacterial strain. It was shown that the mechanism of such dependence is related to the composition and the thickness of the film constituted by corrosion products and formed on the superhydrophobic surface in the course of interaction between the superhydrophobic surface and components of dispersion.

The analysis of the elemental composition of the covering film on the superhydrophobic surface indicated a negligible amount of potassium and sodium in the films formed in PBS. In contrast, a significant amount of sodium was found in the surface films grown in bacterial dispersions. Besides, the films grown on the surface contacting the dispersion of *P. Aeruginosa* were characterized by a higher amount of hydrocarbon components, which were seemingly formed as a product of cell metabolism in the alkaline conditions compared to the films on superhydrophobic surfaces contacting the dispersion of *K. pneumoniae*. The formation of biofilms and sodium polyphosphate films provided enhanced barrier properties for magnesium dissolution and hence for dispersion medium alkalization, leading to inhibition of magnesium substrate degradation. Electrochemical data obtained for superhydrophobic samples continuously contacted the corrosive bacterial dispersions during 48 h indicated a high level of protective anti-corrosion properties of fabricated superhydrophobic coatings and their ability to partially inhibit the medium alkalization. Corrosion currents for the samples immersed in bacterial dispersions in phosphate-buffered saline, followed by drying in an oven had the values of the order of 10^-10^ – 10^-11^A/cm^2^. These currents are 5-6 order of magnitude lower than the currents characteristic of the bare alloy which did not contact to corrosive medium.

## Acknowledgments

The work was supported by the Russian Foundation for Basic Research (grant #18-29-05008) and by the Ministry of Science and Higher Education of the Russian Federation. The XRD measurements were performed using the equipment of CKP FMI IPCE RAS; the SEM and EDX studies were performed using the equipment of the IGIC RAS Joint Research Centre for Physical Methods of Research.

